# Double-layered two-directional somatopleural cell migration during chicken body wall development revealed with local fluorescent tissue labeling

**DOI:** 10.1101/2021.10.28.466242

**Authors:** Nobuyuki Sakamoto, Hirohiko Aoyama, Koji Ikegami

## Abstract

The ventral body wall is derived from the somatic layer of the lateral plate mesoderm, somatopleure, and somite. The primary ventral body wall is formed as a result of the lateral growth of the somatopleure. The secondary body wall is generated through the migration of somitic cells into the somatopleure. While it is reported that the cervical somatopleural cells migrate caudally to the thoracic region during body wall development, the migration of the thoracic somatopleural cells has not been elucidated. To investigate the migration behavior of the somatopleural cells in the thorax during chicken ventral body wall development, we labeled the thoracic somatopleural cells of one somite wide by DiI labeling or gene transfection of enhanced green fluorescent protein and observed the three-dimensional distribution of the labeled cells with the tissue-clearing technique FRUIT. Our labeling experiments revealed the rostral migration of the somatopleural cells into a deep part of the thoracic body wall in embryonic day 6.5 chickens. For embryonic day 8.5 chickens, these deep migrating somatopleural cells were found around the sternal ribs. Thus, we identified the double-layered two-directional migrating pathways of the somatopleural cells: the rostral migration of the deep somatopleural cells and the lateral migration of the superficial somatopleural cells. Our findings imply that the rostral migration of deep somatopleural cells and the lateral migration of superficial ones might be associated with the developing sternal ribs and the innervation of the thoracic cutaneous nerves, respectively.

**Mini-abstract:** Double-layered two-directional migrations of the somatopleural cells in the thoracic body wall during chicken development were revealed by one-somite-wide labeling of the somatopleure.

## Introduction

The ventral body wall, which encloses the visceral organs, is developed in two steps. First, the somatic layer of the lateral plate mesoderm, the somatopleure, extends laterally and joins at the ventral midline to form the primary body wall. Subsequently, muscle and rib primordia derived from the somite migrate laterally into the primary body wall to form the secondary body wall (Nichol et al. 2011; Nichol et al. 2012). In amniotes, the ribs, especially the lower ribs, are slightly bent rostrally toward the sternum (Scaal 2021). This bending is particularly pronounced in birds (Scaal 2021). The ribs of birds comprise of two compartments; the vertebral ribs and the sternal ribs, which are the medial and lateral parts of ribs, respectively, and the ribs bend significantly toward the sternum at the boundary of these two parts (Aoyama et al. 2005). Thus, the rib primordial cells migrate not only laterally but also toward the sternum during the thoracic body wall morphogenesis (Mekonen et al. 2015; Fogel et al. 2017).

Previous studies have shown that the somatopleure is essential for the development of chicken sternal ribs. Sudo et al. showed that the somatopleure-derived inductive signals mediated by BMP are required for the development of sternal ribs through ectopic expression of Noggin, an antagonist of BMP (Sudo et al. 2001). The ectopic limb generated by transplanting the prospective limb somatopleural mesoderm in the thoracic region leads to defects in sternal ribs, indicating that the presence of the somatopleure of the appropriate rostrocaudal level around the sternal rib primordia is required for normal development of the sternal rib (Liem and Aoyama 2009). In rib development, the sternal rib primordial cells migrate into the somatopleural mesoderm and form the sternal rib in the thoracic somatopleure (Burke and Nowicki 2003; Nowicki et al. 2003). Thus, the somatopleure has an important role in the development of the sternal rib.

In the avian embryo, the developmental origin of various tissues of the ventral body wall has been investigated using quail-chick chimera (Chevallier 1975; Chevallier 1977; Aoyama and Asamoto 2000; Shearman et al. 2011). Recently, Hirasawa et al. showed on transplantation of quail somatopleure into chicken, the cervical somatopleure moves to the thorax during avian development (Hirasawa et al. 2016; Nagashima et al. 2016). Furthermore, it was shown that when the somatopleural cells at the level of the forelimb are labeled with the fluorescent dye DiI, the labeled cells were distributed in the sternum (Bickley and Logan 2014). These studies have revealed the dynamic migration of the cervical somatopleure during ventral body wall development through lineage trace experiments.

Previously, we have studied the migration of somitic cells during development of the ventral body wall (Aoyama et al. 2005; Liem and Aoyama 2009). While the importance of the somatopleure in the development of the sternal rib is understood, the origin and migrating pathway of the somatopleure around the sternal rib have not been sufficiently investigated. Given that the somatopleure interacts with the somite during their development, somatopleural and somitic cells may migrate in the same way during thoracic body wall development. To understand the role of the somatopleure in the development of the ribs and thoracic ventral body wall, we need to elucidate the migrating behavior of somatopleural cells of the inter-limb region in thoracic ventral body wall development. Local labeling of the thoracic somatopleure will uncover the migration of somatopleural cells during ventral body wall development.

Recently, various tissue clearing techniques have been well-developed for the imaging of deep tissues, especially to visualize neurons in neurology (Richardson and Lichtman 2015; Tian et al. 2021). The tissue clearing techniques are capable of visualizing the three-dimensional (3D) distribution of cells at the single-cell level through advanced fluorescence microscopy, which allowed imaging clear tissues in chicken embryonic brains (Gómez-Gaviro et al. 2017), and the technique has been used for 3D imaging of the development of GnRH neurons in human fetal brains (Casoni et al. 2016). In chicken development, the migration of neural crest cells (Morrison et al. 2017) and 3D distribution of parasympathetic neurons have been revealed by using the tissue clearing technique (Watanabe et al. 2018).

The closure of the ventral body wall is very important in development. Its failure can cause serious ventral body wall defects (VBWD), such as gastroschisis, bladder exstrophy, and ectopia cordis, in humans (Torres et al. 2015). Though the cause of these congenital anomalies is unclear, Sadler suggested that failure to fold the somatopleural mesoderm may be the cause (Sadler 2010). It is proposed that the movement force of folding is provided by the ventral migration of somitic cells (Sadler 2010). Conditional deletion of Wntless, which encodes the receptor of Wnt ligands, during development of the somatopleural mesoderm causes defects in the migration of somatopleural cells and ectopia cordis in mice (Snowball et al. 2015). Six4/5 expressed in the somatopleure is involved in the closure of the ventral body wall in mice (Takahashi et al. 2018). Fibloblast growth factors expressed in the somatopleure are essential for the ventral body wall formation in mice (Boylan et al. 2020). In addition, double KO mice of TGFß2/3, which are expressed in the somatopleure, also show VBWD (Dünker and Krieglstein 2002). Inhibition of Rho-associated kinase signaling, which is involved in developmental processes, such as cell migration and invasion, causes VBWD in chickens (Riento and Ridley 2003; Duess et al. 2016). Investigating the migrating behavior of somatopleural cells may contribute toward understanding the mechanisms of VBWD in amniotes.

In the present study, we shed light on the migration of thoracic somatopleural cells during the morphogenesis of ventral body wall formation by labeling the small region of somatopleure and analyzing the 3D distribution of labeled cells in the thoracic ventral body wall. Our results revealed the double-layered two-directional migrating pathways of the thoracic somatopleural cells during development of the thoracic ventral body wall.

## Materials and Methods

### Embryos

Fertilized eggs of White Leghorn chickens (*Gallus gallus domesticus*) were purchased from a local farm. The chicken eggs were incubated at 38°C in a humidified incubator and staged according to Hamburger and Hamilton (HH) (Hamburger and Hamilton 1951).

### DiI labeling and 3D reconstruction

CM-DiI (Molecular Probes) at 1 mg/mL in 100% EtOH was diluted to 0.5 mg/mL with 3 M sucrose solution. To label the somatopleure, DiI solution was injected between the ectoderm and somatopleure using a pulled glass capillary (GD-1, Narishige, Tokyo, Japan) by mouth pipetting. The embryos were incubated until the desired developmental stages and fixed using 4% paraformaldehyde in phosphate-buffered saline (PBS) at 4°C overnight. For cryosectioning, the fixed embryos were immersed in PBS containing 30% (w/v) sucrose, and then embedded in Tissue-Tec O.C.T. compound (Sakura Finetek, Tokyo, Japan). The sections (14 μm thick) were prepared with cryostat (CM 3050, Leica Biosystems Nussloch, Nußloch, Germany) and observed under a fluorescence microscope with a UPlanFL N 40**×** objective lens (NA 0.75) (OLYMPUS, Tokyo, Japan). Images were taken with a digital camera (EOS kiss X7i, Canon, Tokyo, Japan) and processed using Fiji (ImageJ; National Institutes of Health) software. The labeled cells, ribs, and nerves were traced manually on GIMP (v2.10, open-source software, http://www.gimp.org/) software and reconstructed with the Fiji 3D viewer plugin.

### Plasmids

Tol2-enhanced green fluorescent protein (EGFP)-C1 was prepared by digesting pT2-7xTcf-NLS-CNL-CP (a gift from Yasushi Okada (Addgene plasmid #65715), Takai et al. 2015) with MluI and inserting PCR-amplified EGFP from pEGFP-C1 (Clontech Laboratories, Mountain View, CA, USA). Tol2-mCherry was produced by digesting Tol2-EGFP-C1 with NheI/EcoRI to remove GFP and inserting mCherry amplified by PCR from 8xGliBS-IVS2-mCherry-NLS-polyA-Tol2 (a gift from James Chen (Addgene plasmid #84604), Mich et al. 2014). pCAGGS-T2TP was gifted by Sato and Takahashi (Sato et al. 2007).

### Electroporation

A plasmid mixture (1.0 µg/µL each of Tol2-EGFP-C1 and pCAGGS-T2TP, 0.4% Fast green or 1.0 µg/µL each of Tol2-mCherry and pCAGGS-T2TP, 0.4% Fast green) was injected into the 21-25 somite level of the coelom using a pulled glass capillary (GD-1, Narishige, Tokyo, Japan) by mouth pipetting. Transfection was performed with an electroporator (CUY21EX, BEX, Tokyo, Japan) under the following conditions: one cycle of a poring pulse of 40 V, on-time 10 msec, off-time 10 msec, and five cycles of a driving pulse of 4 V, on-time 10 msec, off-time 10 msec. The transfection procedure was slightly modified from a previously described method (Fan et al. 2018).

### Tissue clearing

After adequate incubation, the transfected embryos were fixed using 4% paraformaldehyde in PBS at 4°C overnight. After fixation, embryos were washed three times in PBS for 1 h. For tissue clearing, the FRUIT method was used (Hou et al. 2015). The fixed embryos were incubated in 40% FRUIT solution at 37°C for 12 h, and in 80% FRUIT solution at 37°C for 24 h. The embryos were dissected and visceral organs were removed after incubation in 40% FRUIT solution before immersing them in 80% FRUIT solution as necessary. Images of transfected cells in cleared embryos were taken as a z-stack image using an AxioZoom.V16 microscope equipped with a PlanNeoFluar Z 1.0**×** objective lens (NA 0.25) and an ApoTome.2 (Zeiss, Jena, Germany), and processed with Zen2 pro software (Zeiss, Jena, Germany) or Volocity software (Quorum Technologies Inc., CA, USA).

## Results

### The migration of somatopleural cells in the thoracic ventral body wall as revealed by DiI local labeling

To clarify the migration of somatopleural cells during ventral body wall morphogenesis, we labeled the right somatopleure of the 23rd somite level at embryonic day (E) 2.5 (HH 14-15) of chickens by injecting the lipophilic dye DiI between the somatopleure and ectoderm (Fig. 1a, a’). The position and size of the labeled cells were restricted in the pocket between the somatopleure and ectoderm. We expected that labeling a small area could help trace the migration of somatopleural cells. The labeled embryos were incubated until E 7.5 (HH 31-32) and fixed. The thoracic body wall of the labeled embryo was sectioned, and examined for the distribution of DiI-labeled cells (Fig. 1b). DiI-positive cells were found mainly in the lateral region of the thoracic ventral body wall in transverse sections (Fig. 1c, d). In the section at the fifth sternal rib level, DiI-positive cells were found around the sternal rib in the deep layer of the body wall (Fig. 1c). In the section at the fifth vertebral rib level, DiI-positive cells were found in the subcutaneous superficial layer of the body wall (Fig. 1d). Thus, two populations of DiI-positive cells found in different layers of the body wall were distributed at different rostrocaudal levels, and the DiI-positive cells around the sternal rib were in a relatively rostral position compared to the subcutaneous DiI-positive cells (Fig. 1b, c, d).

**Fig. 1.**
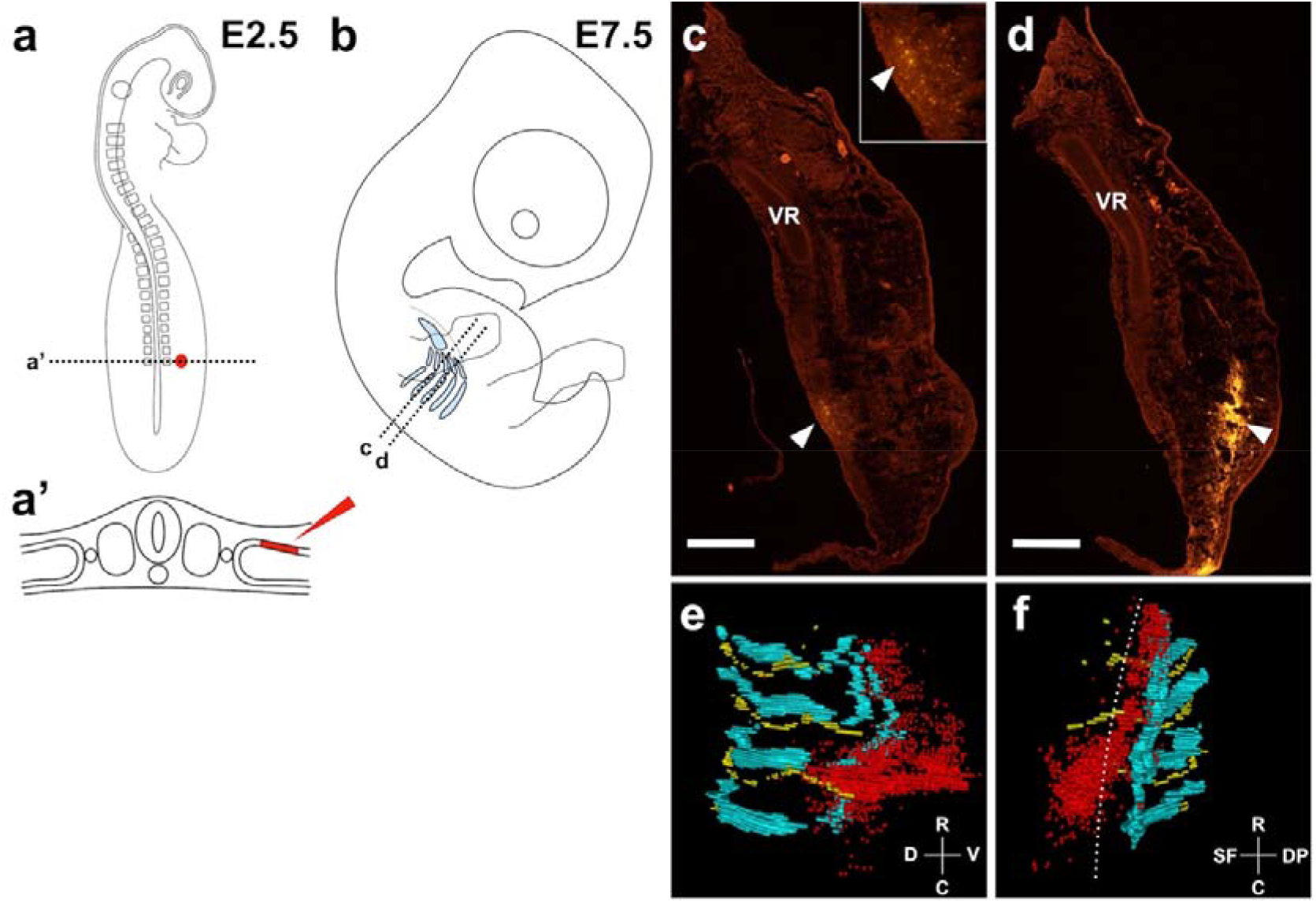
Distribution of the somatopleural mesoderm labeled by DiI. **a** Schematic diagram of a chick embryo at embryonic day (E) 2.5 (HH stage 15) showing the DiI injection site (red point). **a’** Schematic of the transverse section at dotted line a’ in **a**, showing the DiI-labeled domain of the somatopleure (red). **b** Schematic diagram of a chick embryo at E 7.5 (HH stage 31). The ribs and sternum are shown (light blue). Lines c, d indicate the levels of the section shown in **c** and **d** in this plate, respectively. **c** Transverse section of line c in **b**. DiI-labeled cells (white arrowhead) are in the deep layer of the body wall. The inset is a higher-magnification image of the labeled area. **d** Transverse section of line d in **b**. DiI-labeled cells (white arrowhead) are in the superficial layer of the body wall. **e, f** Three-dimensional reconstruction of DiI-labeled cells (red), ribs (light blue), and nerves (yellow). **e** Lateral view, **f** Ventral view. The dotted line indicates the boundary between superficial and deep layers of the body wall. The cross lines indicate the orientation of the views (**e, f**). VR, vertebral rib; R, rostral; C, caudal; D, dorsal; V, ventral; SF, superficial; DP, deep. Scale bars 500 µm (**c, d**)

To analyze a whole image of the distribution of DiI-positive cells in the thoracic ventral body wall, we created 3D reconstructed images from the serial sections where we marked DiI-positive cells, ribs, and nerves on computer images of the sections. The 3D reconstructed images showed that the DiI-positive cells were mainly located around the fifth rib in the thoracic body wall (Fig. 1e). The superficial population of DiI-positive cells was found over and lateral to the fifth vertebral rib, and the deep population of DiI-positive cells was found around the fifth sternal rib (Fig. 1e, f). The 3D reconstruction image revealed that the superficial population of DiI-positive cells was spread laterally, while the deeper population was spread rostrally along the fifth sternal rib (Fig. 1e, f). Figure 1e shows the superficial population of DiI-positive cells along the cutaneous nerve. Together, a majority of the somatopleural cells migrate in the superficial layer laterally along the vertebral rib in a striped pattern, and some cells migrate rostrally in the deep layer along the sternal rib toward the sternum during thoracic ventral body wall development.

### Local genetic labeling of somatopleural cells and observation of deep tissue using a tissue clearing technique

To better analyze the distribution of labeled cells in the thoracic body wall, we employed a tissue clearing technique along with an optical microscope capable of capturing z-stack images. In the uncleared embryo, the DiI-labeled cells in deep tissue could not be clearly identified (Fig. 2a, a’). To observe the distribution of the transfected cells in the deep layer of the thoracic ventral body wall, we used the FRUIT method for tissue clearing (Fig. 2a, b). The whole E4.5 chicken embryo treated with the FRUIT method was almost transparent, albeit the treated embryo was slightly more swollen than the untreated embryo (Hou 2015, Fig. 2a, b). In the cleared embryo tissue, the DiI-labeled cells in the deep tissue were more clearly identified than those in the uncleared embryo (Fig. 2a’, b’). Thus, this tissue clearing technique enabled us to identify each fluorescent-labeled cell in the deep layer of the thoracic ventral body wall without the need for sectioning.

**Fig. 2.**
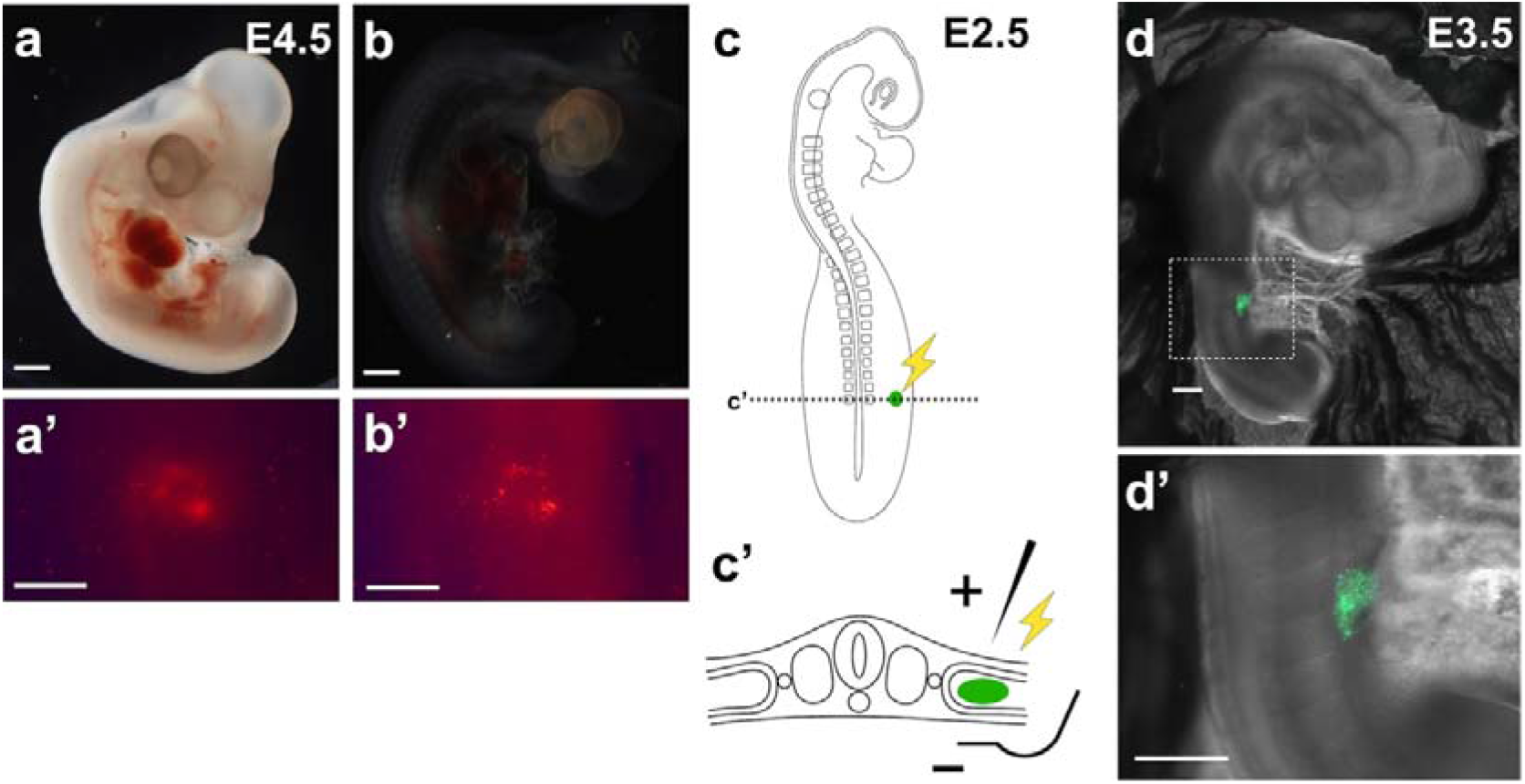
Tissue clearing of the DiI-labeled embryo and the somatopleural mesoderm labeled by EGFP transfection. **a, b** DiI-labeled embryo (E 4.5) before (**a**) and after (**b**) clearing using the FRUIT method. **a’, b’** DiI-labeled cells in **a** and **b. c** Schematic diagram of a chick embryo at E 2.5 (HH stage 15) showing the EGFP transfection site (green point). **c’** Schema of the transverse section at dotted line c’ in **c**, showing the DNA solution in coelom (green). An anode was placed above the dorsal side of the embryo, and a cathode was placed under the embryo. **d** EGFP expression of the transfected embryo (E 3.5) one day after electroporation. **d’** Higher-magnification image of the dotted area in **d**. Scale bars 500 µm (**a’, b’, d, d’**), 1 mm (**a, b**).

For further analysis, we performed labeling of somatopleural cells by another approach. In the above DiI-labeling method, we used the DiI solution at a high concentration to overcome the dilution of the fluorescence following cell proliferation during development. The use of highly concentrated DiI solution might affect the migration behavior of somatopleural cells. In addition, injecting DiI between the somatopleure and ectoderm causes undesired labeling of ectodermal cells, which could generate false results. To avoid these disadvantages, we performed genomic labeling of somatopleural cells using EGFP and transposase expression vectors (Fan et al. 2018). The EGFP gene was electroporated into the somatopleure in E 2.5 (HH 14-15) chicken embryos after injecting plasmids into the coelom (Fig. 2c, c’). Labeling from inside the somatopleure prevents mislabeling of the ectodermal cells. To determine the localization of the labeled cells after gene transfection, we checked the labeled cells one day after labeling. The EGFP-positive cells were observed in the somatopleure within about one somite wide in E 3.5 (HH 19-20) chicken embryos (Fig. 2d, d’). This result demonstrated that genomic labeling with electroporation can label the somatopleural cells in a small area, similar to DiI labeling.

### The aligned migration of deep somatopleural cells in the sternal region of the thoracic body wall

We traced somatopleural cells migrating along the rostrocaudal axis during ventral body wall development. The somite-derived sternal rib primordia penetrated the somatopleure toward the sternum while maintaining their original rostrocaudal position (Chevallier 1975). The somatopleural cells around the sternal ribs may also maintain the original rostrocaudal positions during migration. To confirm this point, we performed genetic labeling by transfection EGFP gene into the various rostrocaudal levels of the somatopleure at E 2.5 (Fig. 3a). Six days after transfection, we examined the distribution of the EGFP-positive cells around the sternal ribs. The transfected embryos were dissected and we observed the labeled cells in the thoracic body wall from their internal surface. The EGFP-positive cells observed at the somatopleure at the 21st, 22nd, and 23rd somite level in the E 3.5 embryos were distributed around the sternal rib of the third, fourth, and fifth rib, respectively, in the E 8.5 embryos (Fig. 3c, d, e). The somatopleural cells between the 24th somite and the rostral half of the 25th somite level in the E 3.5 embryo were distributed around the sternal rib of the sixth and seventh ribs (Fig. 3a, f). The somatopleural cells between the 24th and 25th somite level in the E 3.5 embryo, were distributed around the sternal region of the seventh ribs (Fig. 3a, g). Together, the EGFP-positive cells were distributed around the sternal rib derived from the somite corresponding to the transfected level in the thoracic body wall. Thus, the alignment of somatopleural cells was similar to that when they were transfected during development of the thoracic ventral body wall (Fig. 3a, a’). However, some overlapping distribution of the EGFP-positive cells was observed in the intercostal region, particularly the third and fourth sternal ribs (Fig. 3a’, c, d). Interestingly, the EGFP-positive cells were also in the sternum in a stripe pattern (Fig. 3c, e, f). Thus, the somatopleural cells move toward the sternum, generally maintaining their original rostrocaudal position along the corresponding sternal ribs and forming the ventral body wall.

**Fig. 3.**
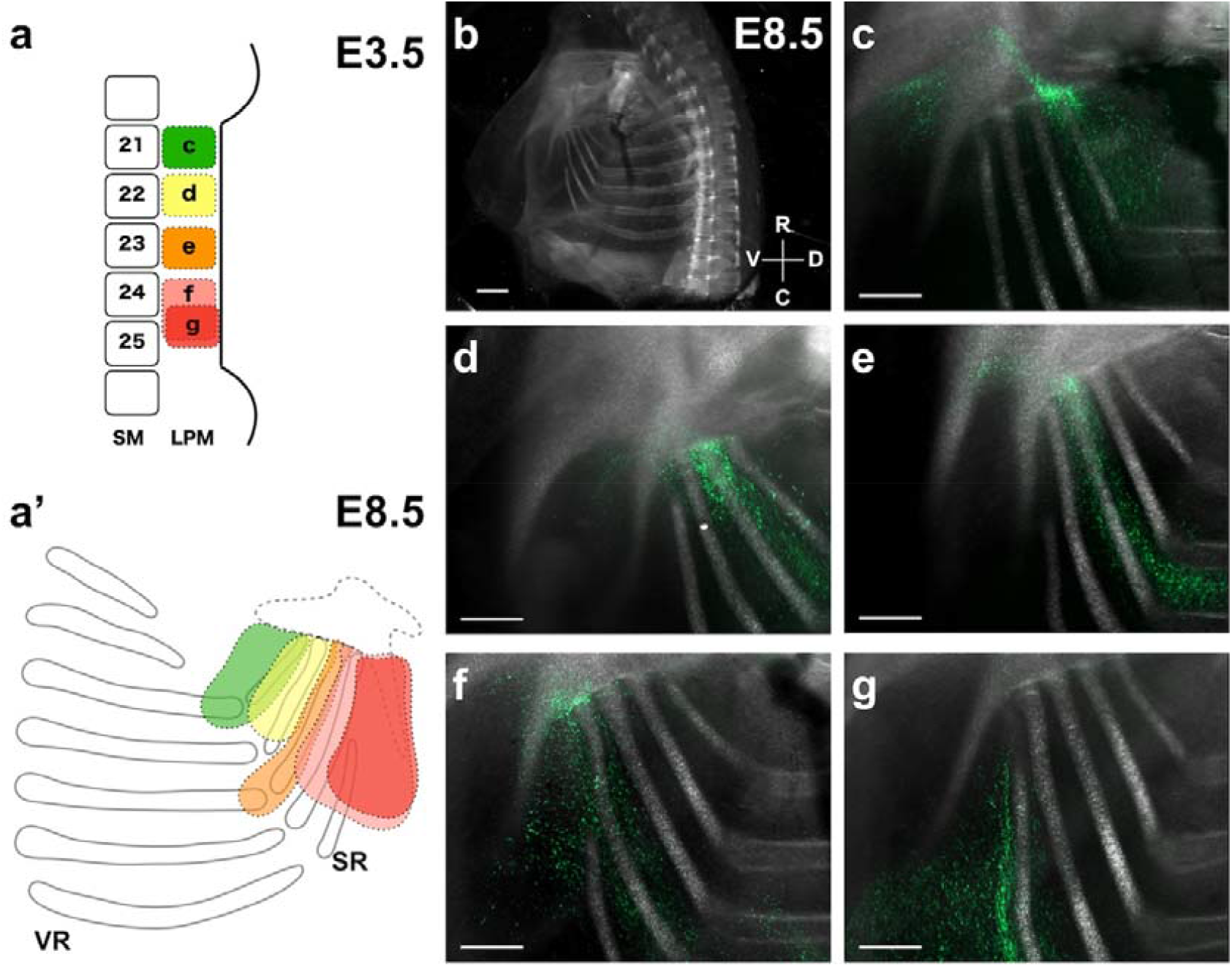
The somatopleural mesoderm migrates in a striped pattern in the sternal region. **a** Schematic diagram of the transfection sites of the embryos (**c–g**). **a’** Schema of migrating regions of the transfected somatopleural mesoderm in the E 8.5 embryo (HH stage 35). The colors indicate the transfected area in **a. b** Medial view of the right thoracic region of the embryo at E 8.5. The cross lines indicate the orientation of the view. **c–g** EGFP-positive cells in the thoracic region of the body wall. SM, somite; LPM, lateral plate mesoderm; R, rostral; C, caudal; D, dorsal; V, ventral. Scale bars 500 µm (**c–g**), 1 mm (**b**)

### The rostral migration of the somatopleural cells during thoracic ventral body wall development as revealed by local genetic labeling using EGFP

We investigated the migration of the somatopleural cells in the chicken embryos from E 4.5 to E 6.5. EGFP and transposase genes were transfected into the right somatopleural cells at the 23rd somite level at E 2.5 (HH 14-15) (Fig. 2a). After 2–4 days of incubation, the right thoracic ventral body walls of the embryos were observed from the lateral side using a fluorescence microscope (Fig. 4a, c, e). In transfected E 4.5 (HH 24-25) embryos, the EGFP-positive cells were distributed in the thoracic ventral body wall with lateral and slightly caudalward elongation compared to E 3.5 (HH 19-20) (Fig. 2b’, 3b). To clarify the 3D distribution of the labeled cells, we analyzed the EGFP-positive cells on the vertical optical yz-plane section images. In the medial section, the EGFP-positive cells were distributed similarly along the rostrocaudal axis from the superficial (left in Fig. 4b’) to deep (right in Fig. 4b’) layer of the body wall. In the middle and lateral sections, the distribution pattern of the EGFP-positive cells was similar to that shown in the medial section, from the superficial (left in Fig. 4b’’ and Fig. 4b’’’) to deep (right in Fig. 4b’’ and Fig. 4b’’’) layer of the body wall. Thus, these yz-plane sections revealed that the EGFP-positive cells migrate laterally in a rectangular form (Fig. 4b’-b’’’).

**Fig. 4.**
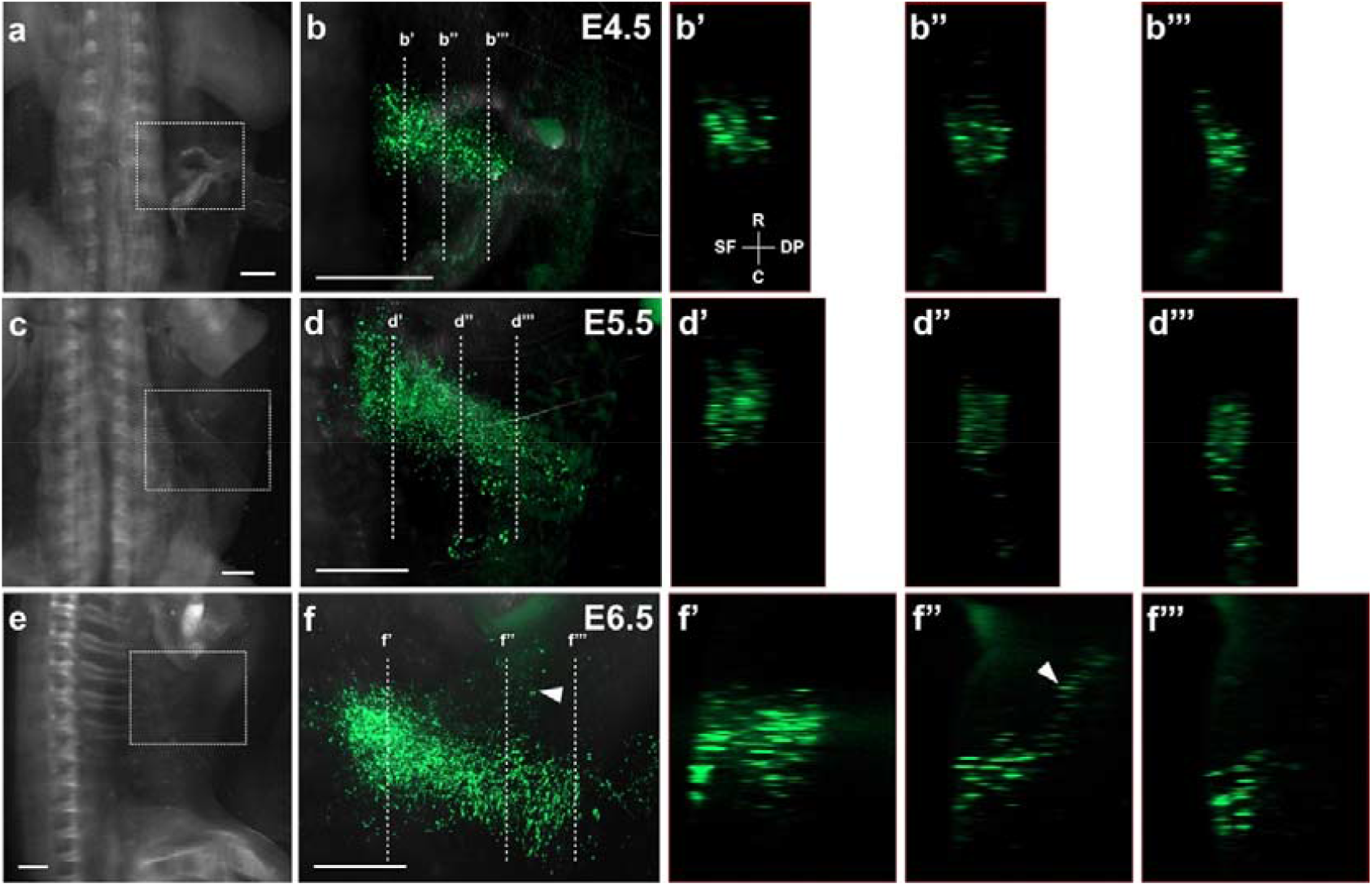
Lateral and rostral migration of the somatopleural mesoderm labeled with EGFP. **a, c** Dorsal views of transfected embryos at E 4.5 (**a**) and E 5.5 (**c**). **e** Lateral view of transfected embryo at E 6.5. **b, d, f** z-stack projections of the dotted area in **a, c**, and **e**, respectively. The dotted line indicates positions of the yz-axis sections (**b’, b’’, b’’’, d’, d’’, d’’’, f’, f’’, f’’’**). The cross line indicates the orientation of the yz-axis sections (**b’)**. White arrowheads indicate the rostral migrating somatopleural cells (**f, f’’**). R, rostral; C, caudal; SF, superficial; DP, deep. Scale bars 500 µm

In transfected E 5.5 (HH 27-28) embryos, the distribution area of EGFP-positive cells extended laterally and slightly expanded rostrocaudally compared with the distribution observed in E 4.5 embryos (Fig. 4b, d). In the optical yz-plane sections of the medial EGFP-positive region in E 5.5 embryos, the transfected cells were distributed with similar rostrocaudal width from the superficial (left in Fig. 4d’) to deep (right in Fig. 4d’) layer of the body wall. In the middle and lateral sections of the E 5.5 embryos, EGFP-positive cells were expanded more rostrocaudally than cells in the medial section (Fig. 4d’’, d’’’). Thus, the distribution of EGFP-positive cells in E 5.5 embryos was similar to that of E 4.5 embryos.

In E 6.5 (HH 29-30) embryos, the labeled cells spread rostralward from the lateral migrating population of the EGFP-positive cells around the joint between vertebral and sternal rib (Fig. 4f, arrowhead). This rostral migrating EGFP-positive cells was spreading from the superficial cell mass to a deeper layer of the ventral body wall in the yz-plane section (Fig. 4f, f’’). Thus, our findings suggest that the somatopleural cells spread laterally first and then a portion of somatopleural cells spread rostrally and deeply. These results coincide with that of the superficial and deep migration pattern of DiI-labeled cells in the ventral body wall (Fig. 1).

### The double-layered distributions of the somatopleural cells derived from different rostrocaudal levels in the sternal region of the thoracic body wall

We investigated whether the somatopleural cell populations that originated from different rostrocaudal levels would settle at the same rostrocaudal level in the secondary thoracic body wall. In E 6.5 (HH 29-30) embryos, some somatopleural cells migrated rostrally into deeper layers of the ventral body wall. Superficial to these rostrally migrating cells, there were other populations of the somatopleural cells that might have been derived from other rostro-caudal levels. Thus, the somatopleural cells derived from different rostrocaudal levels may layer in the ventral body wall. To clarify this point, we electroporated EGFP or the mCherry gene into the different rostrocaudal level of somatopleure. Transfecting two different genes into adjacent regions may cause the double-labeling of the somatopleural cells, because the plasmid mixture was injected into the coelom. To avoid double-labeling, we transfected two different fluorescent genes into the E 2.5 embryos at about one somite wide gap, 22nd–23rd and 24th–25th somite levels (Fig. 5a).

**Fig. 5.**
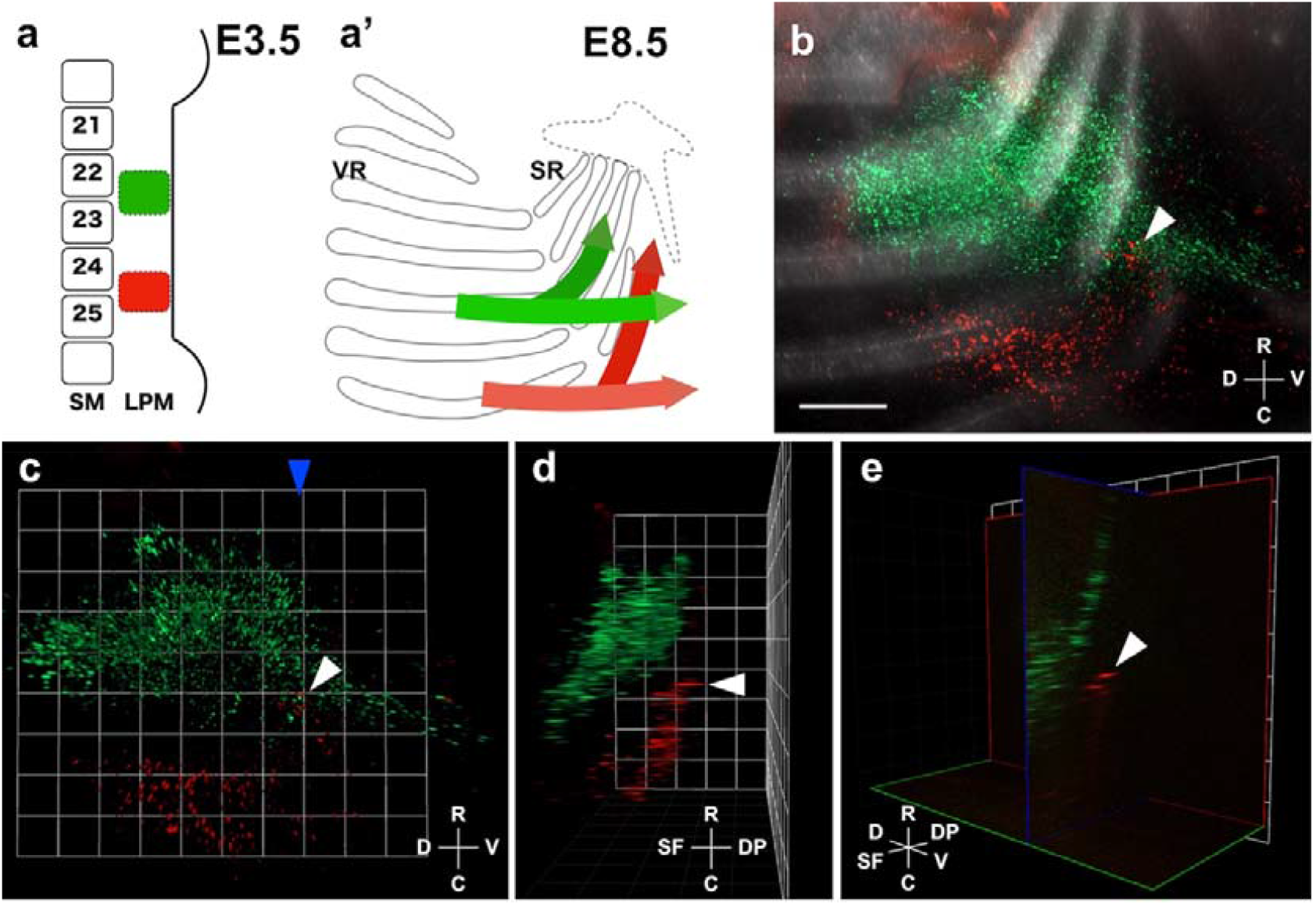
The double-layered distributions of the somatopleural cells derived from different rostrocaudal levels. **a** Schematic diagram of the EGFP (green) and mCherry (red) transfection site in a chick embryo at E 2.5 (HH stage 15). **a’** Schema of two directional migrations of somatopleural cells in the E 8.5 embryo (HH stage 35). Light green and light red arrows indicate the migrating directions of the superficial somatopleural cells. Dark green and dark red arrows indicate the migrating directions of the deep somatopleural cells. **b** z-stack projections of transfected cells in the thoracic body wall. Lateral view. **c** Lateral view of the 3D reconstructed image with Volocity software. The blue arrowhead indicates the position of the yz-slice in **e**. The white arrowhead indicates the area where the two cell populations overlap (**b, c**). **d** Ventral view of the 3D reconstructed image with Volocity software. The caudal somatopleural cells (red, white arrowhead) are beneath the rostral somatopleural cells (green). **e** yz-slice view of the 3D reconstructed image with Volocity software. The white arrowhead indicates the rostral migrating somatopleural cells. The cross lines indicate the orientation of the view (**b, c, d, f**). SM, somite; LPM, lateral plate mesoderm; R, rostral; C, caudal; D, dorsal; V, ventral; L, lateral; M, medial; SF, superficial; DP, deep. Scale bar 500 µm

In the embryos transfected with EGFP and mCherry genes, EGFP-positive and mCherry-positive cells were found at the same area in the sternal rib region in the thoracic body wall at E 8.5 (HH 35) when observed from the lateral side (Fig. 5b, arrowhead). Thus, two cell populations, derived from different rostrocaudal levels of somatopleure, migrated to the same area of the thoracic ventral body wall. To identify the layer of the thoracic body wall, in which these two cell populations are located, we performed 3D analysis (Fig. 5c, d). The frontal view of the 3D reconstruction image and the vertical optical section showed that mCherry-positive cells, which were derived from caudal somatopleure, were in a deeper layer of the ventral body wall compared to EGFP-positive cells, which were derived from rostral somatopleure (Fig. 5d, e, arrowhead). Thus, the deep mCherry-positive cells derived from the caudal thoracic somatopleure spread beneath the superficial EGFP-positive cells derived from the rostral thoracic somatopleure (Fig. 5a’). Together, our findings indicate that the ventral body wall consists of two layers with different rostrocaudal origins.

## Discussion

In chickens, the migration of somitic cells into the somatopleure during this process has been well-studied (Sudo et al. 2001; Nowicki et al. 2003). The migration of cervical somatopleure has been studied in detail (Hirasawa et al. 2016; Nagashima et al. 2016), but the migration of thoracic somatopleural cells has not been sufficiently elucidated. Since the somatopleure interacts with the somite during the development of ribs (Sudo et al. 2001; Liem and Aoyama 2009), we decided to investigate the migrating behavior of the somatopleural cells in the thoracic body wall development. Here, we showed that the somatopleural cells migrate in two directions: the superficial and deep pathways, in chickens. The superficial somatopleural cells migrate laterally and the deep somatopleural cells migrate rostrally along the sternal ribs to form the thoracic ventral body wall. Our findings indicate that the thoracic ventral body wall is composed of two layers of cells of different rostrocaudal origins.

Hirasawa et al. reported that the deep population of cervical somatopleure shifted caudally to the thorax in chickens (Hirasawa et al. 2016). On the other hand, we showed that the deep population of thoracic somatopleure underwent a rostral migration to the sternum, suggesting that the drastic movement of a deep population of somatopleural cells to the thorax, i.e., the sternum along the rostrocaudal axis, occurs during ventral body wall formation. In contrast, the cells distributed in the superficial layer of the somatopleure did not undergo such a rostrocaudal shift. Since somitic cells, which are rib primordia, migrate into the deeper layer of the somatopleure, the somitic cells might be involved in this rearrangement of deep somatopleural cells along the rostrocaudal axis.

We found that the somatopleure at the 21st–25th somite level is distributed in the sternum (Fig. 3). However, Bickley and Logan (2014) showed that the somatopleure at the 14th–21st somite level is distributed in the sternum in their DiI-labeled fate mapping experiments. This discrepancy may be explained by a comparison of the detailed distribution pattern of the labeled cells in the sternum. The somatopleure at the level of the forelimb was distributed in the midline of the sternum, i.e., the sternal keel (Bickley and Logan 2014), whereas the somatopleure at the level of the flanking region was distributed mainly in the lateral part of the sternum, which is close to the ribs (Fig. 3). Thus, the rostral somatopleure migrates caudally and develops into the sternal keel. The caudal somatopleure migrates rostrally and develops into the lateral part of the sternum. These different developmental origins of the skeletal elements of the sternum and different direction of migration of the sternal primordia might indicate the complex and drastic morphogenesis of the avian sternum.

In this study, we showed that some somatopleural cells migrate rostrally and are distributed around the sternal ribs maintaining their original rostrocaudal position (Fig. 3). The sternal ribs are skeletal elements extending from the vertebral portion of the ribs to the sternum in birds and are derived from abaxial somitic cells that migrate into the somatopleure (Burke and Nowicki 2003; Nowicki et al. 2003). The somatopleural cells were shown to migrate in the same area as the primordial cells of sternal ribs that are derived from the same rostrocaudal levels of the somatopleure. This implies that sternal rib primordial cells migrate together with the surrounding somatopleural cells. The interaction between deep somatopleural cells and primordial cells of the sternal ribs might be involved in the migration of somitic cells, which develop into the sternal ribs and are involved in ventral body wall development.

The mechanisms of superficial and deep somatopleural cell differentiation were not elucidated in this study. The rostrally migrating somatopleural cells appear between E 5.5 and E 6.5 (Fig. 4). On the other hand, the abaxial somitic cells, which develop into the sternal rib, appear early at E 4.5 (Nowicki et al. 2003). If the interaction between the somatopleural and abaxial somitic cells induces rostral migrating somatopleural cells, the differentiation of superficial and deep somatopleural cells might occur from E 4.5 to E 6.5.

The direction of migration of superficial somatopleural cells is consistent with the direction of the folding movement of the ventral body wall (Sadler 2010). Therefore, the lateral migration of superficial somatopleural cells may provide a force for the folding movement of the ventral body wall and may be associated with the closure of the ventral body wall. The deep somatopleural cells migrated rostrally to the sternum (Fig. 4, 5). This migrating direction of the deep somatopleural cells may not be consistent with the direction of force for folding movement of the ventral body wall provided by the somitic cells, which migrate ventrally into the somatopleure, as suggested by Sadler (2010). However, since the sternum is derived from the deep somatopleural cells (Fig. 3), the rostral migration of deep somatopleural cells may be involved in the development of the sternum and may be associated with VBWD, such as ectopia cordis. From this perspective, the two directional migrations may be responsible for different causes of VBWD. There is currently insufficient information available on the mechanism of the migration of the somatopleure in the formation of the secondary body wall. Further studies are needed to elucidate the role of the somatopleure in ventral body wall formation.

During the formation of the ventral body wall, first, the somatopleure extends laterally. The superficial somatopleural cells distribute laterally along the cutaneous nerves (Fig. 1e). The laterally migrated somatopleural cells locate to the superficial layer of the ventral body wall at both the vertebral and sternal region (Fig. 5). This striped distribution of the superficial somatopleural cells reminded us of the dermatome. The dermis of the ventral body wall is derived from the somatopleure (Christ et al. 1983) and is innervated by cutaneous nerves. The lateral migration of superficial somatopleural cells might be involved in the innervation of thoracic cutaneous nerves. It was reported that BMP4 expressed in the subcutaneous mesoderm is required for the development of the cutaneous nerves of hind limbs (Honig et al. 2005). The dermis of the dorsal body wall is derived from the somite (Olivera-Martinez et al. 2000) and is innervated by cutaneous nerves. The dermatome, which is the result of the innervation of cutaneous nerves, might be formed by the migration of somitic and somatopleural mesodermal cells.

Our present findings suggest that the lateral migration of superficial somatopleural cells and the rostral migration of deep ones might be associated with the innervation of the thoracic cutaneous nerves and the developing sternal ribs, respectively.

## Acknowledgments

We thank the following people for kindly providing the plasmids: Y. Sato and Y. Takahashi for pCAGGS-T2TP, Y. Okada for pT2-7xTcf-NLS-CNL-CP (Addgene plasmid #65715), and J. Chen for 8xGliBS-IVS2-mCherry-NLS-polyA-Tol2 (Addgene plasmid #84604). Part of this study was supported by JSPS KAKENHI, grant number JP26460254. Part of this work was carried out at the Natural Science Center for Basic Research and Development, Hiroshima University.

